# Quantifying the Prevalence of Non-B DNA Motifs as a Marker of Non-B Burden in Cancer using NBBC

**DOI:** 10.1101/2024.01.04.574106

**Authors:** Qi Xu, Jeanne Kowalski

## Abstract

Alternative DNA structures, such as Z-DNA, G-quadruplexes, and mirror repeats, have shown potential involvement in cancer etiology. NBBC (Non-B DNA Burden in Cancer) is a web-based tool designed for quantifying and analyzing non-B DNA motifs within a cancer context. Herein, we provide a step-by-step protocol for employing NBBC, starting with data input and proceeding through the quantification and normalization of non-B DNA motifs that result in calculation of non-B burden. With detailed instructions for performing various motif-centric analyses based on cancer gene signatures, including DNA damage repair and response pathways for exploring genomic stability, and sample-level gene mutation signatures, a user is able to explore non-B associative correlations within current cancer biology. We provide additional detail on input queries into NBBC, interpret the quantitative results, and apply normalization techniques to ensure accurate comparisons across different genomic regions and non-B DNA structures.

NBBC offers a powerful and user-friendly interface for the cancer research community. This chapter serves as an essential, enhanced instructional guide for researchers to leverage NBBC in their cancer biomarker investigations for an understanding of the potential role of non-B DNA in contributing to them.

## 1 Introduction

Non-canonical DNA refers to DNA structures that differ from the canonical B-DNA double helix structure, including G-quadruplexes, cruciform, slipped structures, triplexes, and Z-DNA(1-4). It has been discovered that non-B DNA-forming sequences can induce genetic instability in human cancer genomes, suggesting a role in cancer development (1).

While there are several non-B DNA databases and prediction tools that exist, the majority of these tools primarily focus on individual motif sequences in isolation (5,6). We introduce the concept of “non-B Burden” as a quantitative marker, to provide the capacity to integrate information from non-B DNA motifs into a comprehensive, genome-wide perspective. This viewpoint has been notably absent in prior non-B DNA research, which underscores its innovative nature and potential. A parallel concept in cancer research can be found in the idea of tumor mutation burden. As tumor mutation burden quantifies the prevalence of mutations and can inform on cancer prognosis and treatment response, our introduction of “Non-B Burden” holds a similar promise for assessing non-B DNA motif prevalence and its potential for interpretation of biological processes, particularly within the realm of cancer research.

To accelerate the non-B burden analysis in cancer, NBBC is designed to serve as a new analysis and visualization platform for the exploration of non-B DNA. NBBC serves as a valuable resource for researchers investigating the role of non-B DNA structures in cancer and other genetic diseases.

In this chapter, we demonstrate a ‘how to’ guide to access and explore NBBC (7)for gene signature analyses that may be further combined with familiar downstream correlatives. In doing so, we present details on the quantitative approach and normalization methods that are available at various genomic levels, including the gene-level, signature-level, and sample-level. Altogether, we provide a guided approach to the use of NBBC, including input query, quantification, and normalization of non-B DNA motifs, and how to use non-B burden in downstream applications.

## 2 Materials

### 2.1 The web server, NBBC

To simplify the use of non-B burden calculation and introduce it for wide, non-bioinformatic research uses, we introduce NBBC, A Non-B DNA Burden Explorer in Cancer. NBBC is an online web server that provide non-B burden calculation, non-B burden visualization and non-B motif exploration. NBBC serves to conduct non-B burden computations and offers normalizations that enable comparisons across genes or non-B structures. It provides visualizations for descriptive analysis of burden values, burden distribution, and burden-based gene clustering. The NBBC webserver is accessible without any login requirements and is completely free to use (https://kowalski-labapps.dellmed.utexas.edu/NBBC/).

### 2.2 Non-B DNA Motif Data

The Non-B DNA forming motif data in NBBC are downloaded from Non-B DB 2.0 database with hg19 build (5,6,8). There are 7 non-B structure motifs included: A-phased repeat (APR, n = 2,386 motifs), G-quadruplexes (G4, n = 361,232 motifs), Z-DNA (n = 404,192 motifs), inverted repeats (IR, n = 5, 771,570 motifs), mirror repeats (MR, n = 1,378,864 motifs), direct repeats (DR, n = 1,113,354 motifs), and short tandem repeats (STR, n = 2,826,360 motifs). A subset of MR and IR motifs are further delineated within the application to represent Triplex (Triplex-MR, n = 412,028 motifs) and Cruciform (Cruciform-IR, n = 147,152 motifs) motifs, respectively. To ensure the reliability and relevance of our data, we have sourced our non-B DNA data the NBBC web server and so users have no need to download the non-B DNA data.

### 2.3 Query Input

### 2.3.1 Non-B Burden at gene-level

To summarized, The input for NBBC include three major types: a single gene (a list of genes), a gene signature, or genomic coordinates (see below). To accommodate these input types, we have provided four options for users to choose from. In the web server (“Input Page” tab), Option 1 offers pre-populated gene sets related to cancer, while Option 2 provides molecular signatures of cancer cell lines. Alternatively, users may manually input gene symbols in Option 3 or upload genomic coordinates of interesting regions in Option 4.

a. **A single gene**. The typical use case is a quick single gene search by typing the “hgnc symbol” of gene, such as KRAS, BRCA, etc.
b. **Gene signatures**. The input includes popular cancer signatures, cell line molecular signatures or user-defined signatures. NBBC offers built-in cancer-related gene sets for quick query, including cancer hallmark gene signatures from the MSigDB database (9), DNA damage repair and response gene signatures (10,11), and molecular signatures (option 2) from the Genomics of Drug Sensitivity in Cancer database (12).
c. **Genomic coordinates of regions**. This is a general option where the query region is not a full region of a gene but a sub-region of the gene, such as cancer-specific mutation sites or regions with copy number alterations. In this case, user can upload the region in query (genomic coordinates) in a table (see **Note 1**).

#### 2.3.2 Non-B Burden at sample level (Burden in Batch)

Furthermore, NBBC allows users to upload “multiple groups” of genomic regions (genes, mutation regions) in batch and calculate the non-B burden for each group. We named it as “Burden in Batch”. Using “Burden in Batch”, it allows users to calculate the non-B burden for each sample (see **Note 2**) to enable further downstream analysis of associations. The input format of “Burden in Batch” is a table with four columns (*group_id, chromosome, start, end*). Each row represents a genomic region and the “*group_id*” is used to group the genomic regions.

## 3. Methods

### 3.1 How to use NBBC to calculate non-B burden

The NBBC web server provides a user-friendly platform for calculating the non-B DNA burden within genomic sequences. Our protocol illustrates this calculation through two exemplar use cases.

a. **Gene-Level Non-B Burden (Basic Use):** NBBC can process both single and multiple genes to calculate the non-B burden. Here, we utilize the “Homologous Recombination” gene signature (10,11), the maintenance of which plays an important role in cancer genome stability. This illustration aims to demonstrate the foundational computation of non-B burden and emphasize the critical role of normalization techniques in the analysis.
b. **Sample-Level Non-B Burden (Advanced Use)** This second application showcases the “Burden in Batch” feature of NBBC. We employed mutation site data from 511 samples from the TCGA-LGG (Lower Grade Glioma) dataset (13). Each sample comprises a list of mutation sites, complete with chromosomal coordinates. The “Burden in Batch” tool in NBBC is utilized to quantify the number of non-B motifs that overlap with these mutation sites, thereby summarizing the non-B burden for each sample. The objective here is to illustrate the integration of non-B burden analysis with clinical datasets, highlighting its utility in sample-level evaluation.

### 3.2 Start with the “Input Page”

The “Input Page” of the NBBC web server is the starting point for users to input their data for non-B burden analysis. This page is designed to accommodate various user requirements through four distinct input options:

a. **Option 1: Built-In Cancer-Related Gene Signatures** Users can choose from pre-defined gene sets related to cancer, such as those involved in DNA damage repair, response gene pathways, cancer hallmark gene sets, and oncogenes, among others.
b. **Option 2: Cancer Cell Line-Specific Features** This option provides a selection of molecular features specific to cancer cell lines, including mutations and copy number alterations.
c. **Option 3: Manual Gene Input** For users interested in conducting a quick query, this interface allows the manual input of single or multiple genes.
d. **Option 4: Genomic Coordinates Upload** Users can upload genomic coordinates that define regions of interest, such as mutation sites, to assess the non-B burden in the context of mutation-localized regions.

With the integration of these input capabilities, NBBC offers comprehensive coverage for non-B burden calculations at varying levels of genomic detail, from precise mutation sites to expansive gene signatures.

**Fig. 1.**
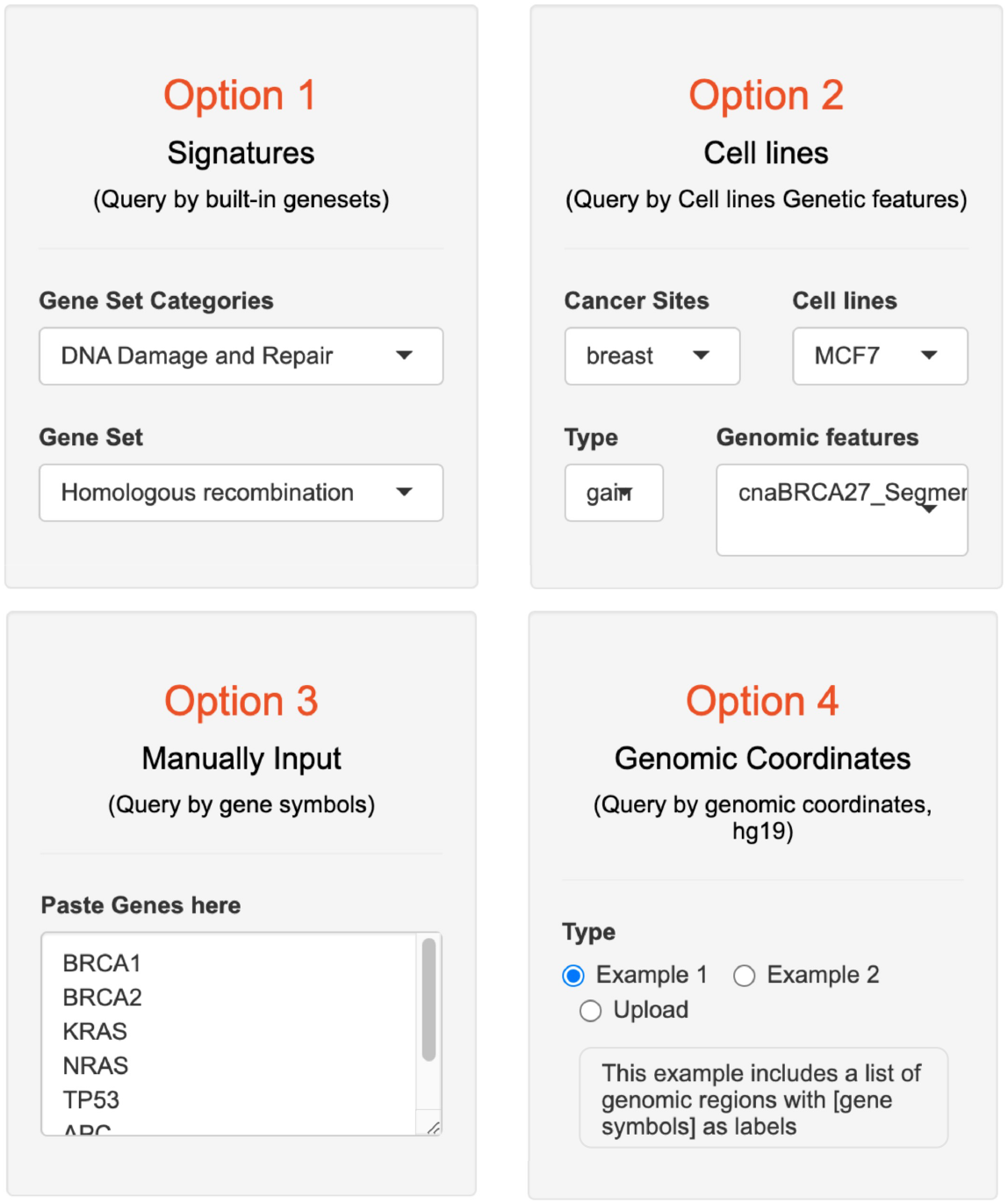
The input page of NBBC. In the “Input Page”, users can select from four distinct options to begin their analysis with NBBC. Option 1 provides a selection of pre-defined gene sets pertinent to cancer research. Option 2 allows users to explore molecular signatures from various cancer cell lines. For a more customized approach, Option 3 enables the manual entry of specific gene symbols. Option 4 permits the uploading of genomic coordinates for regions of interest.

### 3.3 Non-B Burden at gene level (Basic use)

#### 1. Select genes as input

In the input page, under option 1, Select the “Homologous recombination” gene set under the “DNA Damage and Repair” category. There will be a preview window at the bottom to show the selected genes (34 genes). Click “Next > Gene Screen” to navigate to the “Gene Screen” module.

#### 2. The quantification of non-B DNA motif

Non-B Burden is calculated by counting the occurrence of non-B forming regions associated with each specific non-B DNA type. After the query gene list are selected, the non-B burden will be calculated in the background by the web server. And the calculation value will be visualized in bar plots on the right panel of the “Gene Screen” tab page. There are two tabs called “Total Burden” and “Burden by type”.

a. **Total Burden**. NBBC includes a Total Burdens plot that visualizes the total non-B burden in each gene as a bar plot. This visual provides users with a summary of the non-B burdens for each query unit.
b. **Burden by non-B type**. The stacked bar plot shows the non-B burdens by type for each gene in the query. This allows users to easily compare the non-B burdens across different genes and identify genes with potentially higher burdens from certain non-B structures.
c. **Interactive plots**. Since this is an interactive plot, users can select hovers for more details of each data point. This feature allows users to obtain a more detailed understanding of the non-B burden for each gene in the query and can aid in identifying potential targets for further investigation.
d. **Other options**. On the left side panel of this page, there are checkbox options, where NBBC provides users with the option to display only a subset of genes or non-B types. This flexible functionality allows users to tailor their analysis to their specific research needs and enables them to focus on the genes or non-B types of interest.

#### 3. The normalization of non-B burden

Normalization is a crucial step in ensuring meaningful comparisons of Non-B Burden across diverse genes and non-B DNA structure types (see **Note 3**). The NBBC provides several metrics for normalization to facilitate these comparisons.

**Fig. 2.**
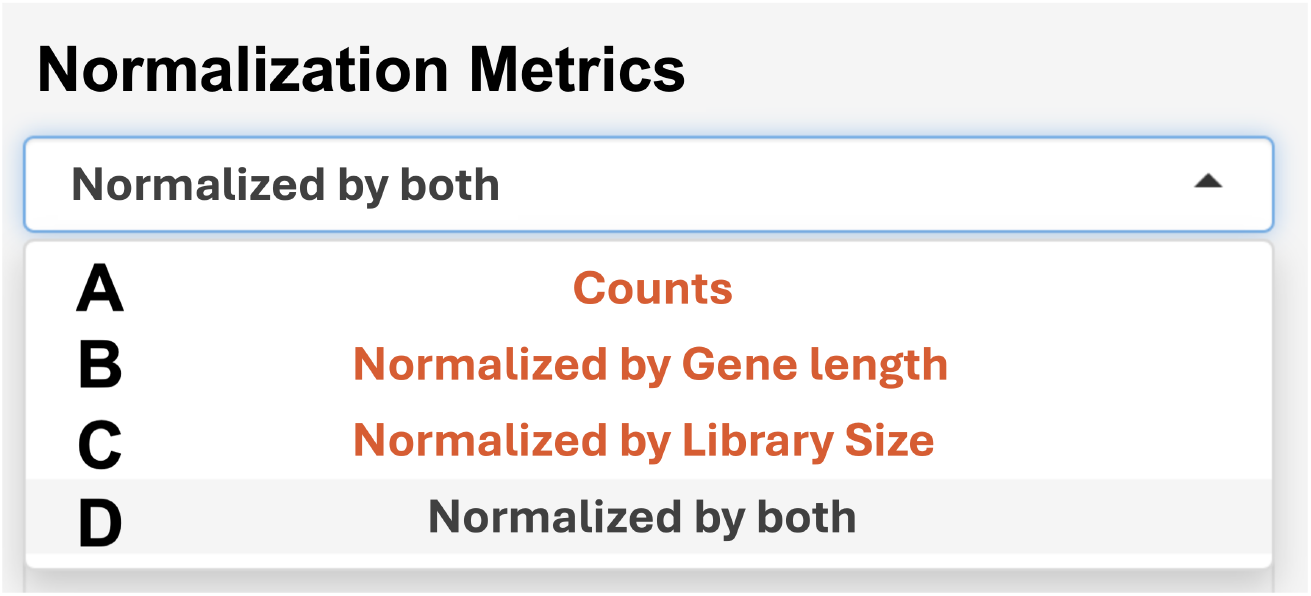
The normalization options for non-B burden. (**A**) “Count” indicates raw motif counts, serving as the unnormalized base measure. (**B**) “By Gene Length” adjusts Non-B Burden based on gene size for comparability across genes. (**C**) “By Library Size” scales Non-B Burden to the non-B motif library size, aiding in analyzing various non-B DNA types. (**D**) “Normalization by Both” offers a measure standardizing Non-B Burden by both gene length and motif library and is the default normalized metric in NBBC.

a. **Raw Motif Counts:** The basic measure of Non-B Burden without any normalization, representing the number of non-B motifs present.
b. **Normalization by Gene Length:** Adjusts the Non-B Burden by the length of the gene region. This is critical when comparing different genes, as it accounts for the varying sizes of genomic regions.
c. **Normalization by Motif Library Size:** Tailors the Non-B Burden relative to the size of the non-B motif library. This normalization is particularly useful when assessing the burden of different types of non-B DNA within a single gene.
d. **Normalization by Both (CPKM):** This comprehensive measure normalizes Non-B Burden both by the length of query regions (per kilobase, 10^3^) and by the library sizes of non-B motifs (per million, 10^6^), facilitating a comparison across both genes and non-B motif types. This default unit, CPMK (counts per kilobase per million), ensures a standardized comparison by considering both the prevalence of non-B motifs and the scale of the genomic and motif libraries.

Normalization allows for the comparison of Non-B Burden across both different genomic regions and among distinct types of non-B DNA for the assessment of differences in motif prevalence among them. Specific applications of normalization include (**Fig.3**):

**Fig. 3.**
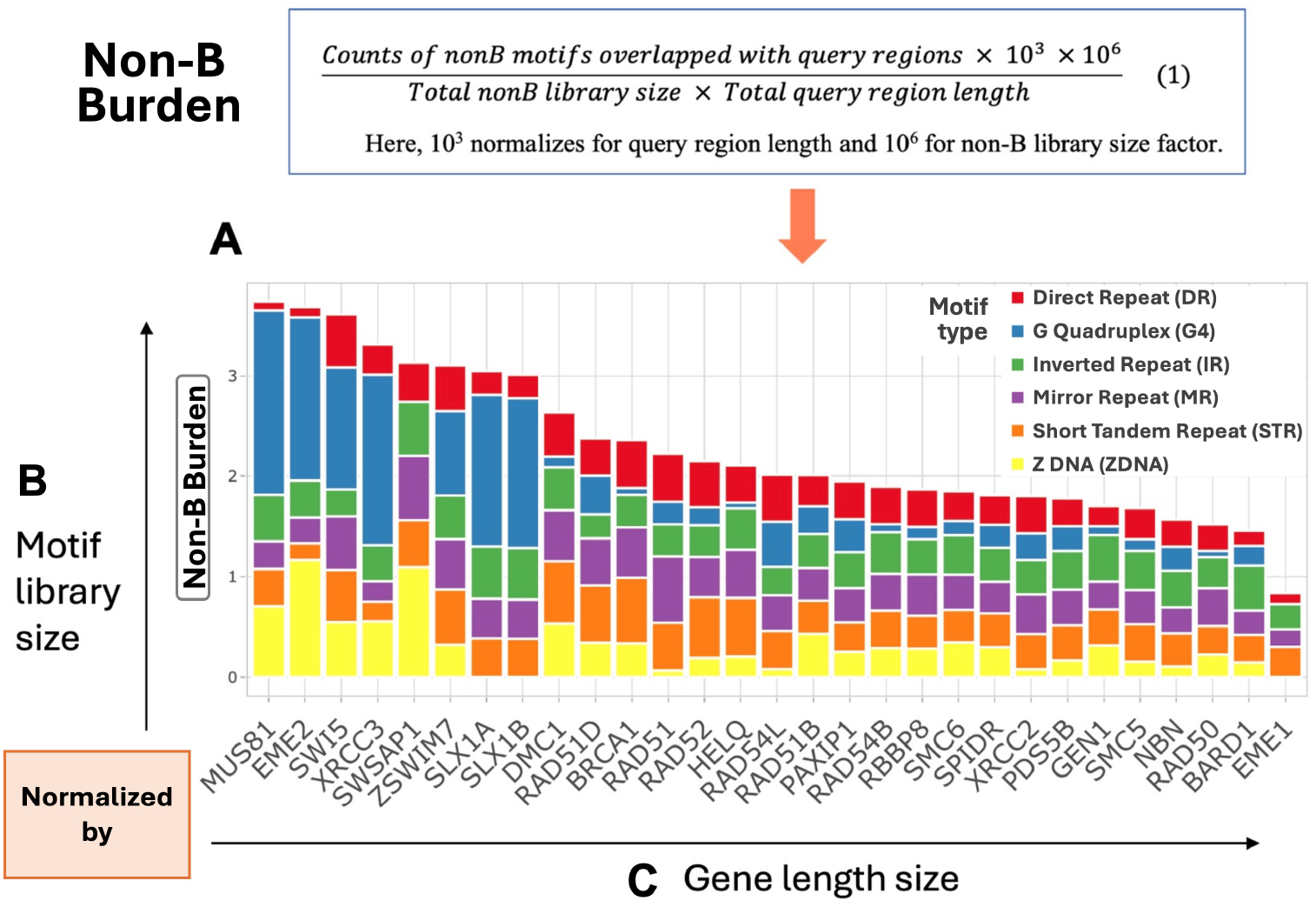
The Non-B Burden distribution in Homologous recombination signature queried in NBBC. **(A)** The non-B burden is visualized in a bar plot with normalization across both genes and non-B types. **(B)** The normalization by non-B motif library sizes allow burden comparison across non-B types. **(C)** The normalization by gene length enables comparisons across genes.

a. **Motif Library Size Normalization:** Recommended when the aim is to compare the Non-B Burden across various non-B DNA types within the same gene. It adjusts for the total number of motifs in each non-B motif type, providing a direct comparison of prevalence across different structures.
b. **Gene Length Normalization:** Advised for comparing Non-B Burden across multiple genes within a signature. This accounts for the potential bias where longer genes might inherently contain more non-B motifs, thus offering a more accurate reflection of non-B motif density, rather than sheer quantity, within the gene’s region.

The aim of this part (**Gene-Level Non-B Burden**) in NBBC is to conduct a gene-level analyses of non-B representation that could prove helpful to focus on a single or subset of genes of interest for hypothesis generation.

### 3.4 Non-B Burden at sample level (Advance use)

#### 1. Objective of “Burden in Batch”

a. **Conception**. The ‘Burden in Batch’ function extends NBBC’s capabilities, enabling the computation of non-B burden at the sample level. It allows for the analysis of multiple groups of genomic regions, such as mutated regions across different tumor samples, facilitating a deeper exploration of non-B burden in relation to clinical data.
b. **Application**. This feature is especially beneficial for generalized queries based on extensive sets of genomic regions. These can represent distinct tumor samples, thereby enabling the calculation of sample-specific non-B burden. This analysis can reveal potential links between non-B burden and clinical outcomes.

This function can be access under the “Burden in Batch” tab from the home page in NBBC (**Fig.4A**).

**Fig. 4.**
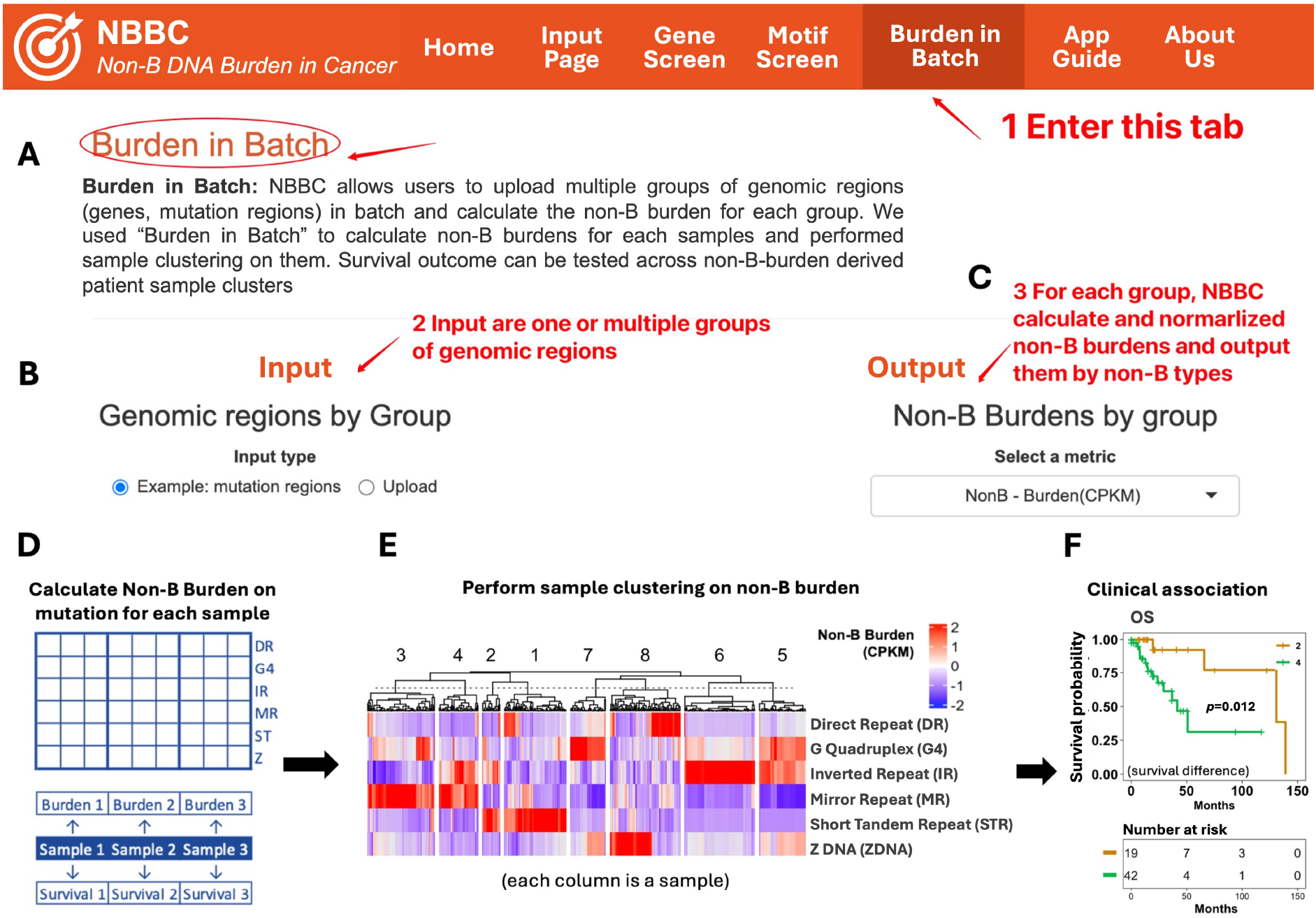
The “burden in batch” analysis in NBBC. **(A)** The web interface to “Burden in Batch” analysis to calculate non-B burden at sample level in batch. (B) The input area to upload the query genomic region grouped by group ids. **(C)** The output area to output non-B burden with normalization options. (D) A graphical summary of non-B burden calculation at the sample level with mapped survival data. **(E)** The heatmap visualizes the clustering results of TCGA-LGG sample based on mutation-localized, sample-level non-B burdens. **(F)** Cluster 1 (STR high) and Cluster 4 (MR-IR high) show a significant overall survival (OS) difference (p = 0.0012).

#### 2. How to perform “Burden in Batch” analysis in NBBC

- **Prepare input table**. The input format of “Burden in Batch” is a table with four columns (*group_id, chromosome, start, end*), —collectively referred to as the “query table”. The example data can be download by clicking the “Download Example” button in the page. Each row represents a genomic region and the “*group_id*” is used to group the genomic regions. The typical use case for “Burden in Batch” is to calculate non-B burden for each tumor sample using a list of genomic coordinates.
- **Case Study Input:** For our example, we calculate mutation-localized non-B burden for TCGA Lower Grade Glioma (LGG) samples (n=511), using genome-wide mutation site regions as the query input. In other words, 511 groups of genomic mutation regions from 511 LGG samples will be use as input for the calculation (see **Note 4**).
- **Calculate non-B burden for each sample**. Users compile a query table containing the genomic mutation regions for all samples, ensuring columns are accurately labeled. Upon navigating to the “Burden in Batch” input page, users upload their table and initiate the analysis(**Fig.4B**). The server quantifies non-B motifs coinciding with the input regions and applies the default CPKM normalization. All the non-B motifs overlapped with mutation regions of each sample will be summarized for non-B burden at sample level. On the right side of the page, a matrix will be output to show non-B burden from different type (columns) of each sample (rows). Users can choose to download the metrics by click the “Download Burden Output” button.
- **Choose an appropriate normalization method**. Similar to the normalization methods gene-level non-B burden, the sample-level non-B burden also support four types of metrics, including “counts”, “normalize by motif library size”, “normalize by motif region length” and “normalize by both (CPKM)”. One difference here is the “region length”. Different “gene length”, the region length here are the total length of all the query regions for each sample (**Fig.4C**).
- **Perform sample clustering on non-B burden**. Subsequent to calculating the non-B burden matrix, we perform clustering analysis on the samples based on their non-B burden. The data is standardized using Z-score normalization, with k-means clustering applied in this instance. But users can choose other preferred clustering methods. The purpose here is to showcase how this sample-level quantification of non-B burden can be used for downstream analysis (**Fig.4E**).
- **Associate clusters with clinical data**. The clustering process categorizes samples into groups characterized by distinct non-B burden profiles. These groups can then be utilized in further analyses, such as survival or association studies, to elucidate the potential clinical significance of non-B burden variations in cancer (**Fig.4F**).

NBBC serves as a valuable resource for researchers investigating the role of non-B DNA structures in cancer and other genetic diseases. By offering an accessible platform for analyzing and visualizing non-B DNA burden within a cancer context, NBBC enables the quantification and exploration of non-B DNA by a wide, non-bioinformatic user base.

## Acknowledgements

The website of NBBC is hosted on an AWS server provided by the Kowalski Lab and is supported by UT-Austin IT solutions and Dell Medical School at UT-Austin.

## Notes

1. Each row in the table represents a region in query. The four column names are required to be the same as the example [*hgnc_symbol, chromosome, start, end*]. [*hgnc_symbol*] can be either gene name or any tags to annotation the region but may not be duplicated. [*chromosome, start, end*] takes genomic coordinates as input.
2. “Burden in Batch” is different from “Genomic coordinates of regions”. Although they both take genomic coordinate as input, “Genomic coordinates of regions” only allows one region under one label, such as one sub-region in a specific gene. So in its input table, the “*hgnc_symbol*” column should be unique. On the other hand, “Burden in Batch” allows the query of a list of regions under one label, such as all the mutated regions in one tumor sample. Therefore, in its input table, the “*group_id*” column (such sample id) will be repeating for different genomic coordinate. Then non-B burden can be summarized for this sample.
3. The concept of normalization in RNA-seq analysis (14), exemplified by metrics like CPM (Counts Per Million), RPKM (Reads Per Kilobase of transcript, per Million mapped reads), plays a similar role as in our approach for normalizing Non-B Burden. Similar to RNA-seq, where these normalization techniques ensure the comparability of gene expression values across diverse samples, the normalization methods employed in non-B Burden calculations serve a similar purpose.
4. The mutation data of TGCA-LGG is download from UCSC Xena (13,15) (https://tcga-xena-hub.s3.us-east-1.amazonaws.com/download/mc3%2FLGG_mc3.txt.gz). The Xena hub is at https://tcga.xenahubs.net and the title is “dataset: somatic mutation (SNP and INDEL) - MC3 public version”.

